# Lower limb in sprinters with larger relative mass but not larger normalized moment of inertia

**DOI:** 10.1101/2022.02.25.482057

**Authors:** Natsuki Sado, Hoshizora Ichinose, Yasuo Kawakami

## Abstract

**Purpose:** Sprinters exhibit inhomogeneous muscularity corresponding to musculoskeletal demand for sprinting execution. An inhomogeneous morphology would affect the mass distribution, which in turn may affect the mechanical difficulty in moving from an inertia perspective; however, the morphological characteristics of sprinters from the inertia perspective have not been examined. Here we show no corresponding differences in the normalized mass and normalized moment of inertia between the sprinters and untrained non-sprinters.

**Methods:** We analyzed fat- and water-separated magnetic resonance images from the lower limbs of 11 male sprinters (100 m best time of 10.44–10.83 s) and 12 untrained non-sprinters. We calculated the inertial properties by identifying the tissue of each voxel and combining the literature values for each tissue density.

**Results:** The lower-limb relative mass was significantly larger in sprinters (18.7 ± 0.7% body mass) than in non-sprinters (17.6 ± 0.6% body mass), while the normalized moment of inertia of the lower limb around the hip in the anatomical position was not significantly different (0.044 ± 0.002 vs. 0.042 ± 0.002 [a. u.]). The thigh relative mass in sprinters (12.9 ± 0.4% body mass) was significantly larger than that in non-sprinters (11.9 ± 0.4% body mass), whereas the shank and foot relative masses were not significantly different.

**Conclusion:** We revealed that the mechanical difficulty in swinging the lower limb is not relatively larger in sprinters in terms of inertia, even though the lower-limb mass is larger, reflecting their muscularity. We provide practical implications that sprinters can train without paying close attention to the increase in lower-limb mass and moment of inertia.

## Introduction

Human morphological characteristics are primarily the result of adaptation to bipedal locomotion (1, 2). Human morphology enables bipedal walking and endurance running efficiency, resulting in current humans being good endurance runners but poor sprinters owing to the lack of a galloping mode (2). Meanwhile, the sprinting velocity in humans (≈10 m/s) is higher than the maximum velocity in the quadrupedal trotting mode, which is biomechanically comparable to human sprinting (3). The sprinting ability in humans exhibits considerable individual variability; for example, the Japan National Record of 50 m sprint is 5.75 s while the average time of 50-m sprint in Japanese males is approximately 7.37±0.52 s (4). This wide individual variability is attributed to a combination of training and genetic factors. The morphology of a well-trained athlete is informative for enhancing the motor performance in humans.

Muscle morphology in athletes has been examined as an indication of potential motor performance, based on the exertion abilities of the joint torque (5) and power (6), which are primarily determined by muscle size, especially muscle volume (5, 7). Previous MRI studies on sprinters have demonstrated greater trunk and lower limb muscles in sprinters than in non-sprinters (8–10). The muscularity was shown to be inhomogeneous with the specific development of the hip flexors and extensors (8–10). Within the same functional groups (quadriceps and hamstrings) of thigh muscles, sprinters exhibit an extremely large semitendinosus and rectus femoris (8, 11). These muscles have maximal cross-sectional areas at more proximal thighs than other muscles (e.g., the semitendinosus compared with the semimembranosus and the rectus femoris compared with the vastus medialis) (12). The muscle sizes of the hip extensors and flexors are related to the running velocity and/or 100 m sprint time in sub-elite (8, 9, 13) and elite sprinters (10). It has also been reported that hip extensors’ volumes discriminate elite sprinters from the sub-elite (10). Muscularity in sprinters corresponds well with the musculoskeletal demands that increasing running velocity in the high-speed range (>7 m/s) requires a larger torque/power exertion of the hip flexors and extensors (14, 15).

From a mechanical perspective, any motion ***a*** :acceleration, ***α*** :angular acceleration,) is determined by the combination of force-related (***f***: forces, ***τ***: torques) and inertia-related (*m*: mass, ***I***: moment of inertia) factors in line with Newton-Euler equations of motion:

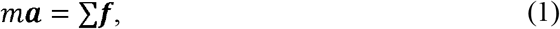

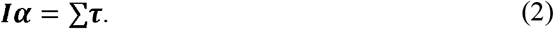

As humans control their motion via joint torques induced by muscle forces, the relationship between the torque and moment of inertia (Eq. 2) is particularly important for human motor execution. The above muscular characteristics in sprinters (8–10, 13) can be interpreted as an difference in force-related factors, but simultaneously, the morphological differences would involve differences in the mass distribution and thereby inertia-properties. As lean tissue density is larger than that of fat tissue (16, 17), larger muscles would lead to a larger lower limb mass, leading to a general speculation of the trade-off between greater torque exertion ability and greater difficulty in moving the lower limb (i.e., moment of inertia). However, the moment of inertia *I* is mechanically proportional to the square of the radius of gyration *r*:

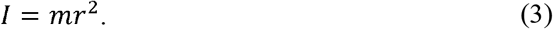

As discussed above, sprinters have particularly large hip flexors and extensors (8–10). Within the thigh muscles, the muscles whose maximal cross-sectional areas are located at the proximal thigh (12) are particularly developed in sprinters (8, 11). Muscular inhomogeneity may lead to a greater mass closer to the hip in sprinters. Owing to the proximal-specific larger muscularity in sprinters, the moment of inertia may not increase linearly with the mass. Anatomists have previously developed scaling coefficients to calculate body-segment inertia parameters (BSIPs) for motion analysis. This is done using the direct measurements of elderly cadavers (18) and indirect measurements *in vivo* through medical images or surface scans such as gamma-ray scanning (19) computed tomography (20), magnetic resonance imaging (MRI) (21, 22) and three-dimensional (3D) laser scanners (23). However, the inertial characteristics of athletes have not been fully examined.

Based on the simple mechanical laws shown in Eq. 2), an increase in torque exertion and a smaller rate or no increase in the moment of inertia would lead to a larger angular acceleration, which could in turn facilitate improved sprint performance. Understanding morphology from an inertial perspective would provide useful knowledge for the development of human motor performance. In this study, we compared the normalized inertia properties of sprinters to those of untrained non-sprinters by analyzing water- and fat-separated MRIs. Among the inertial properties, we focused on the normalized lower-limb moment of inertia around the hip joint 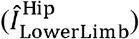 because of the high musculoskeletal demand in the lower limb swing during sprinting (14, 15). We hypothesized that the lower limbs in sprinters have a larger relative mass but not a larger 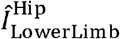 than non-sprinters.

## Methods

### Experimental design

We performed an *a priori* power analysis for an independent *t*-test with the parameters of type I error: *α=*0.05, statistical power: 1-*β=*0.8, and effect size: Cohen’s *d*=1.2 ‘very large’ (24), which calculated the total sample size (*n*) as 24 (*n*=12 in each group). According to the power analysis, we recruited a total of 24 healthy males, including 12 well-trained (athletic career >7 years and 100 m best records: 10.44–10.83 s) sprinters and 12 non-sprinters who did not undertake resistance training or sports activities for at least 2 years. However, the data of one of the sprinters could not be analyzed because of an MRI error (fat/water separation error in the Dixon sequence) around the endpoint of the toe in the foot segment scan, and the main outcome data (i.e., the lower-limb mass and the moment of inertia around the hip joint) could not be calculated. Thus, we excluded this participant’s data from the analysis. The analyzed participants were 11 sprinters (age: 20 ± 2 yrs, height: 1.77 ± 0.06 m; body mass: 68.2 ± 4.4 kg, Mean ± SD) and 12 non-sprinters (age: 23 ± 3 yrs, height: 1.69 ± 0.04 m; body mass: 58.6 ± 4.5 kg). The purpose and experimental protocol of the study were explained to the participants and they provided written informed consent. The Human Research Ethics Committee at Waseda University, Japan approved the study protocol (reference number:2019-174).

We used a 3.0 Tesla MR scanner (SIGNA Premier, GE Healthcare, Milwaukee, USA) to obtain the MR scans. All lower limb segment scans were performed in the supine position. The 2-point 3D-dixon axial sequence, called “Lava-Flex” in GE Healthcare, was performed on the right thigh, shank, and foot segments (‘Spacing-Between-Slices’: 2 mm, ‘Pixel-Spacing’: [0.2930 0.2930] mm, ‘Repetition Time’: 4.3 ms, ‘Echo-Time’: 1.7 ms, ‘Rows’ × ‘Columns’: 1024 × 1024 pixels, other detail information can be confirmed in the Supplemental Table S1). Each scan of the thigh and shank had two sections because of the limitation of the length scanned in one section. We prevented the deformation by placing pads of an appropriate size below the pelvis, knee, and heel, thereby lifting the soft parts of the thigh and shank off the bed. The two sections of each thigh and the shank scans were taken continuously without changing the position on the MRI machine. In the MRI settings, the anteroposterior and mediolateral coordinates of the two consecutive scans coincided. To minimize the effect of magnetic field inhomogeneity on the Dixon sequence, the superoinferior ranges of the two scans overlapped by > 10 cm.

### MRI processing

We analyzed the 16-bit depth MRI images using MATLAB 2019a (MathWorks, Natick, MA, USA). A flowchart of the segmentation procedure is presented in Fig. 1. To create a temporal body mask, we first created fat-only and water-only temporal binary images using a low intensity threshold. These binary images were added together, and the temporal body mask was defined by filling any internal holes in the added masks based on a binary morphology operation (25). After creating the temporal body mask, we removed the pixel intensity in the unwanted areas (other than the range of interest, such as the part of the opposite leg segment included in the image). After that, we estimated and then homogenized the inhomogeneity field using a grayscale morphological closing operation with a large circular structure element (25). The threshold for the foreground and background signal intensity of each of the fat-only and water-only images was determined using Otsu’s method (26), which created water-binary and fat-binary images. We created a definitive body mask by adding the two masks together and filling any internal holes with a binary morphology operation (25). We classified each pixel as a fat tissue, lean (muscle or skin) tissue, or a background using water- and fat-binary images. For a slice having some voxels involved in both water- and fat-binary images, we further performed *k*-means clustering to classify the intensity of the voxels in the sum of the fat-only and water-only images (i.e., in-phase image) into three categories (fat tissue, lean tissue, or background), which were applied to the voxels involved in both the water-binary and fat-binary images. The trabecular bone has MRI intensity characteristics similar to fat tissue because of the yellow marrow (27), while the cortical bone is not brightened by MRI. These bone tissues were classified as follows:

**Figure 1.**
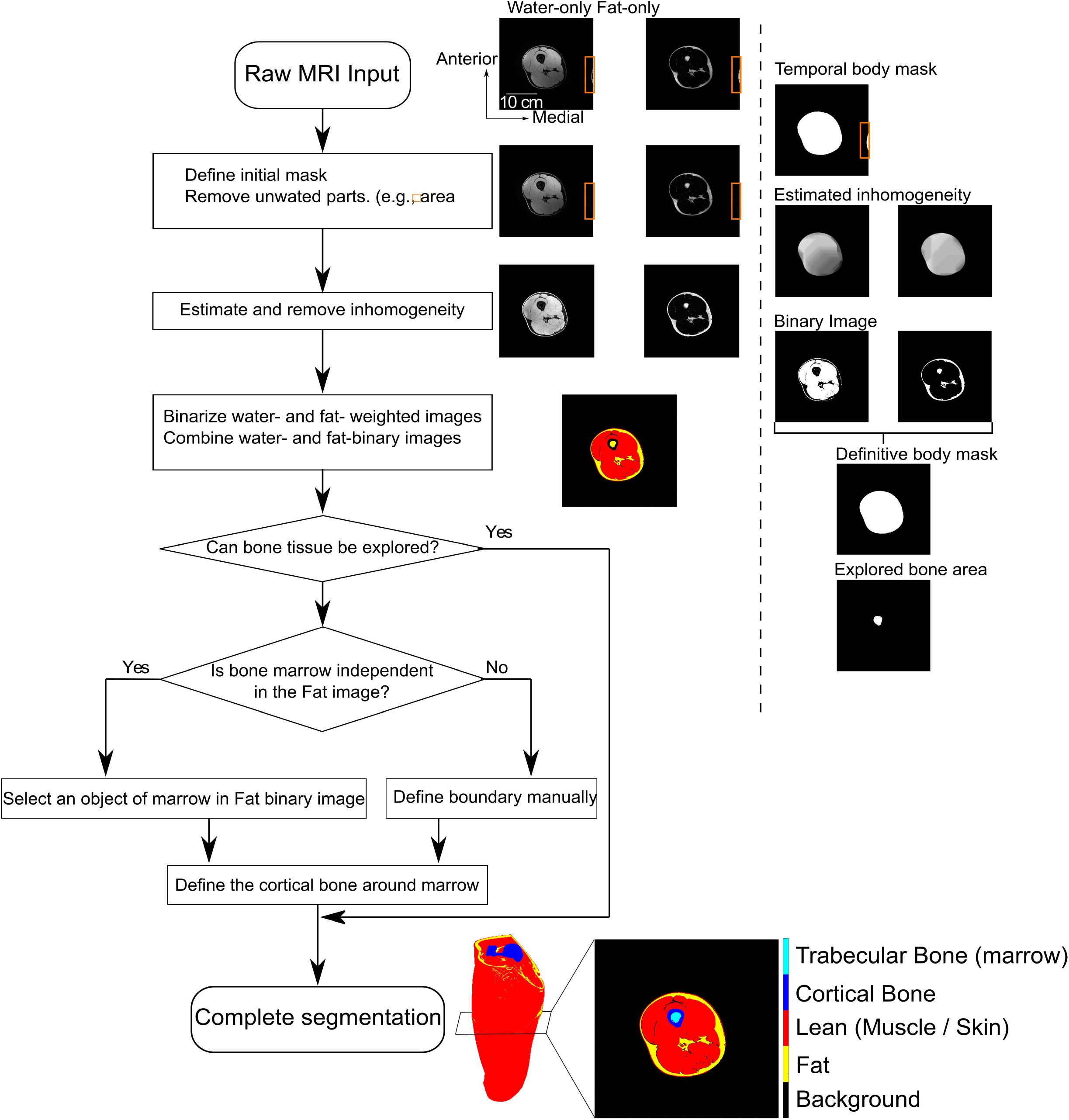
Flowchart of the segmentation process based on water/fat separation images.

> Step 1. Search for a region inside the body mask region where the pixels classified as fat tissue (trabecular bone in fact) are surrounded by pixels classified as background (cortical bone in fact) and define them as trabecular bone and cortical bone.
>
> Step 2. (When a bone cannot be explored in Step 1 and the trabecular bone is an independent object in the fat-binary image), select and define the independent object in the fat-only image as a trabecular bone.
>
> Step 3. (When a trabecular bone was in contact with the outer fat tissue in the fat-binary image and could not be defined in Steps 1 and 2) Manually draw a border with the fat in the fat-only image or in-phase image using a pen tablet (Artist 15.6 Pro, XP-PEN, Japan) and define the object as a trabecular bone.
>
> Step 4. (After Steps 2 and 3) Define the pixels that surround the trabecular bone and are classified as background (cortical bone in fact) as a cortical bone.
>
> Step 5. Adjust the cortical bone if necessary (for example, if the tissue adjacent to the cortical bone that is not depressed on MRI is considered to be a cortical bone).

### Data calculation

The 3D coordinates of the anatomical landmarks (see Supplemental Table S2) on the MR coordinate system were derived via manual digitizing in the 3D mode using the OsiriX software version 13 (Pixmeo, Geneva, Switzerland) by an examiner. The examiner repeated the digitization of all landmarks from eight participants for whom it had been >1 month following the first digitizing process to assess the intra-examiner repeatability of manual digitizing.

The ankle joint in the foot and shank scans and the knee joint in the shank and thigh scans were defined as the midpoints between the malleoli and the lateral and medial articular cleft of the knee, respectively. The hip joint was defined using a least-squares calculation for a sphere fitting of 30 points distributed over the surface of the femoral head, similar to Harrington et al. (28). The ankle and knee boundary planes were defined as the horizontal planes passing those joint locations (Fig. 2A). The hip joint boundary was defined by a plane 37° (29, 30) vertical on the medial to the hip joint and a horizontal plane lateral to the hip joint (31) (Fig. 2A). The lean and fat voxels outside of the border were excluded. The cortical and trabecular bone voxel in that segment was retained even though it was outside the proximal and distal borders. Bone voxels within the segment were retained even if they were outside the boundary, whereas bone voxels in another segment were excluded even if they were inside the boundary.

**Figure 2.**
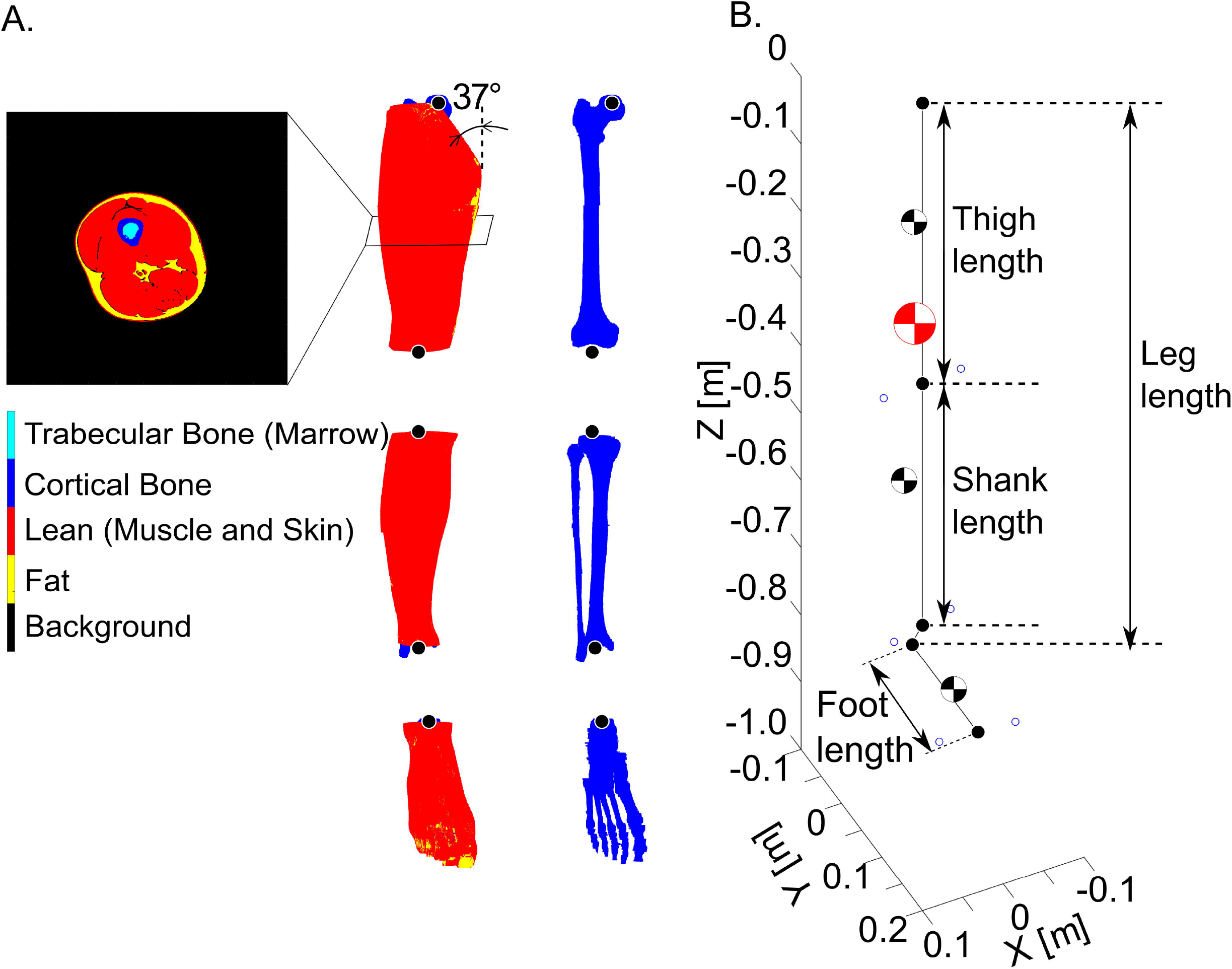
A typical example of the constructed data. A three-dimensional constructed volume data of the thigh, shank and foot (A) and a stick picture created from three-dimensional position coordinates of the entire hypothetically constructed lower limb (B). Note: The origin of the hypothetical coordinates was at the hip joint center. The lower limb length was defined as the height difference between the hypothetical coordinates of the hip and calcaneus.

Based on the literature values (16, 17), we defined the tissue density as:

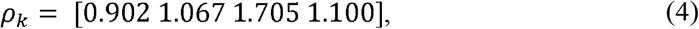

where *k* =1,2,3,4 shows the tissues of fat, lean, cortical bone, and trabecular bone, respectively. The mass, *m*_*s*_, volume *V*_*s*_, density *ρ*_*s*_, and position coordinates of the center of mass (CoM), 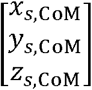, of a segment *s* (*s* = 1,2,3 shows the segments of thigh, shank, and foot, respectively) were calculated as

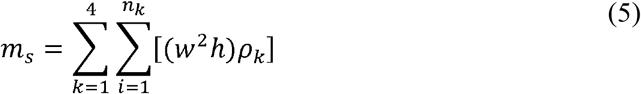

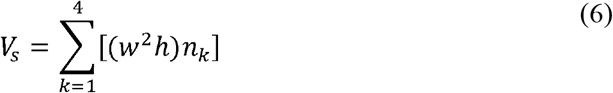

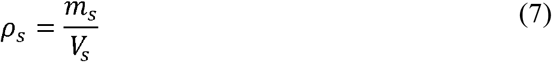

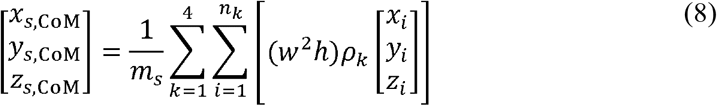

where *n*_*k*_ is the number of voxels classified as the tissue *k, w* and *h* were the width (‘Pixel-Spacing’:0.2930 mm) and height (‘Spacing-Between-Slices’:2.0000 mm) of each voxel (i.e., *w*^2^*h* is the volume of each voxel), and 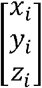 is the position coordinate of the *i*th voxel, respectively.

Inertia tensor of segment *s*, 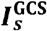, was calculated as shown below:

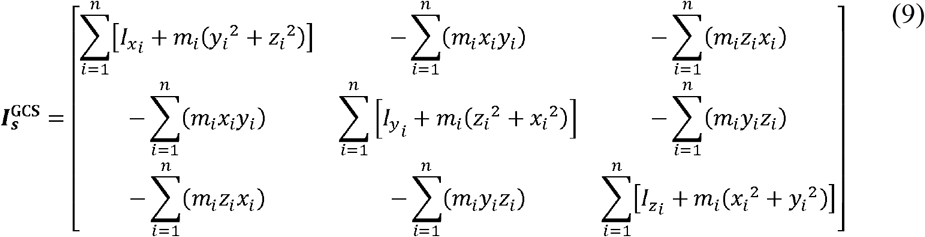

where *m*_*i*_ is the mass of *i*th voxel, 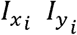 and 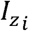 were the moments of inertia of each voxel and were calculated as follows.

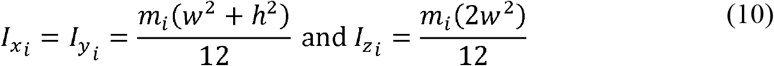

A right-handed orthogonal segment coordinate system (SCS) was fixed at each of the thigh, shank, and foot segments using the 3D coordinates of landmarks. This definition was consistent with that used by Sado et al. (32, 33). We calculated the inertia tensor in each 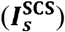 as follows:

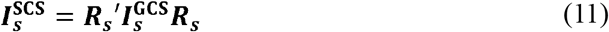

where ***R***_*s*_ is the rotational transformation matrix from the SCS to the GCS of segment *s*. Note that the principal axes of the segments are not consistent with the axes of the SCSs (34); thus, 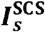 is not in the form of a diagonal matrix, which is similar to Dumas et al. (34), considering the differences between the SCS axes and the inertial principal axes.

We hypothetically created the lower-limb data in an anatomical position from the 3D coordinates in the MRI coordinate system of the thigh, shank, and foot using hierarchically constructed coordinates with the hip as the origin (Fig. 2B). In this hypothetical creation, the ***x***-axis directions of their SCSs were consistent. We calculated the moment of inertia of segment *s* about the extension–flexion (or plantar flexion–dorsiflexion) axis of each proximal joint *j* (*j* = 1,2,3 shows the joints of the hip, knee, and ankle, respectively) as

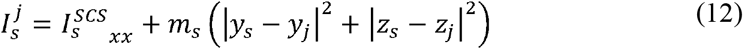

where 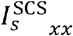 is the moment of inertia of segment *s* around its CoM, *y*_*s*_, *y*_*j*_, *z*_*s*_ and *z*_*j*_ are the anteroposterior or superoinferior coordinates of the CoM of segment ***s*** or joint *j* in a hypothetically created coordinate. We further calculated 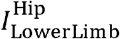 in the data of this hypothetical anatomical position as follows:

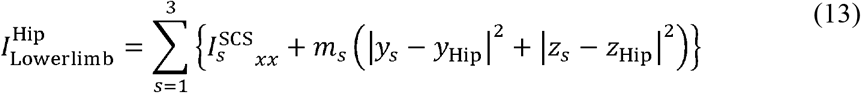

where 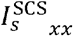 is the moment of inertia of the segment around its CoM, *y*_*s*_, *y*_Hip_, *z*_*s*_ and *z*_Hip_ are the anteroposterior or superoinferior coordinates of the CoM of segment ***s*** or the hip joint in a hypothetically created coordinate. In this coordinate system, the origin is set at the hip joint; thus, *y*_Hip_, and *z*_Hip_ are equal to zero.

The segment and the lower-limb masses were expressed relative to the whole-body mass. Both the products and the moments of inertia of each segment were expressed relative to the segment length *l*_s_ and the segment mass *m*_s_. For example, the moment of inertia of segment *s* around the ***x***-axis is

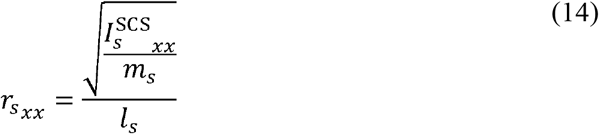

Segment lengths were defined as the Euclidean distances between the proximal anddistal joint locations (Fig. 2B). Similarly, we normalized the moment of inertia of each segment of its proximal joint 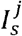.

Using the scaling method established by Hof (35), we normalized 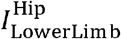 by the lower-limb length (*l*_Lowerlimb_) and the whole-body mass (*m*_Body_) as follows:

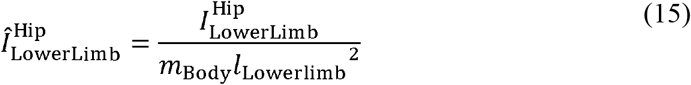

where *l*_Lowerlimb_ is the height difference between the hip joint center and calcaneus in hypothetically created coordinates (Fig. 2B).

### Statistical analysis

We used MATLAB 2019a for statistical analysis. We used intraclass correlation coefficients (ICCs) and coefficients of variation (CVs) to assess the repeatability of the examiners’ manual-digitizing-basis segment lengths. The ICCs can be interpreted as good (>0.75) or excellent (>0.90) (37). After normality confirmation using the Jarque-Bera test, we calculated the Pearson’s correlation coefficient (*r*) to confirm the effect of body size on the lower-limb inertia properties.

We tested the significance of the inter-group differences using an independent *t*-test. The statistical significance was set at 0.05. The effect size of each *t*-test was determined using Cohen’s *d* (36) with classification as small (≤0.49), medium (0.50–0.79), large (0.80–1.19), very large (1.20–1.99), and huge (≥2.00) (24).

## Results

The height and body mass of sprinters were significantly greater than those of non-sprinters (*P* < 0.01, *d* = 1.60, 2.16; Table 1). The absolute values of the lower-limb mass and moment of inertia were significantly correlated with the body size parameters, while the relative mass and 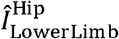 were not (Fig. 3).

**Table 1.**
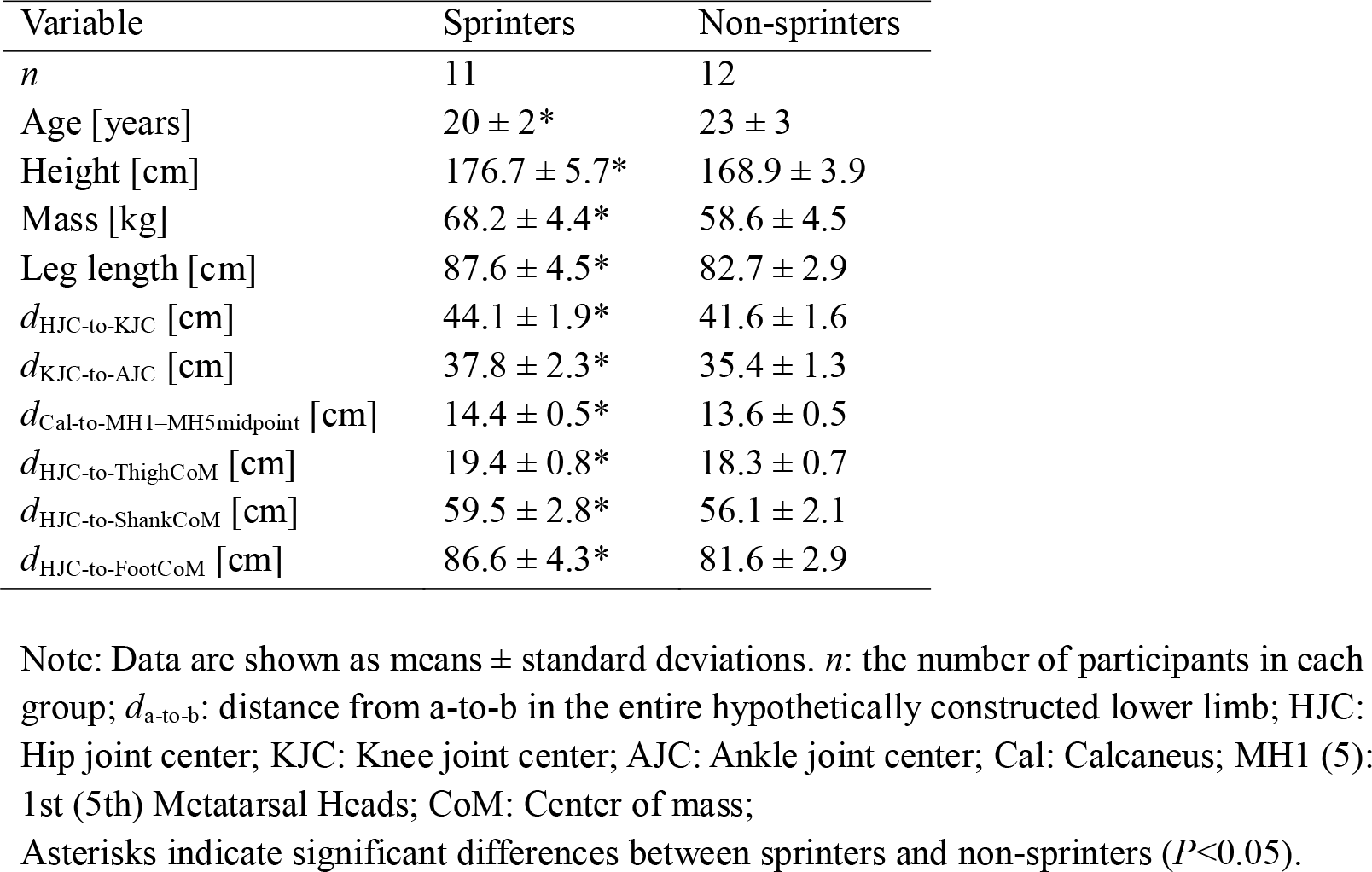
Characteristics of participants.

**Figure 3.**
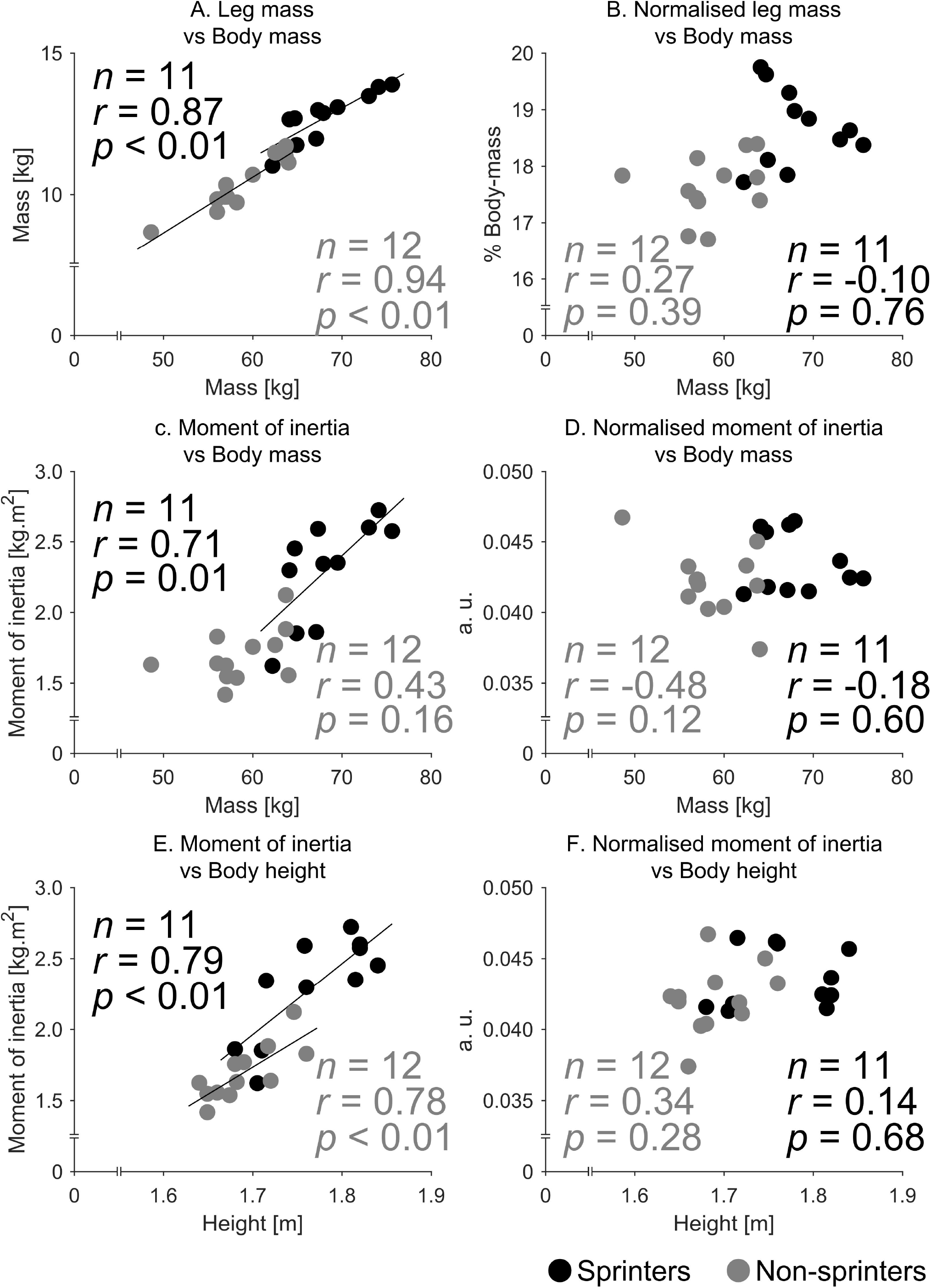
Relationships between absolute (A, C, E: left column) and normalized (B, D, F: right column) inertia parameters and body size parameters (A–D: body mass, E–F: body height) in sprinters (black) and non-sprinters (grey). Note: *n*, and *r* were the sample size and Pearson’s correlation coefficients in each group, respectively.

The relative mass of the entire lower limb was larger in sprinters (18.7 ± 0.7% body mass) than in non-sprinters (17.6 ± 0.6% body mass) (*P* < 0.01, *d* = 1.73; Fig. 4A). However, 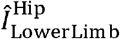 in sprinters (0.044 ± 0.002 [a. u.]) and non-sprinters (0.042 ± 0.002 [a. u.]) did not differ significantly (*P* = 0.15, *d* = 0.62; Fig. 4B). From individual plots, we can observe that, although one of the non-sprinters showed a small 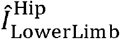 (Fig. 4B), the 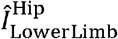 in non-sprinters and sprinters, except for him, were highly comparable.

**Figure 4.**
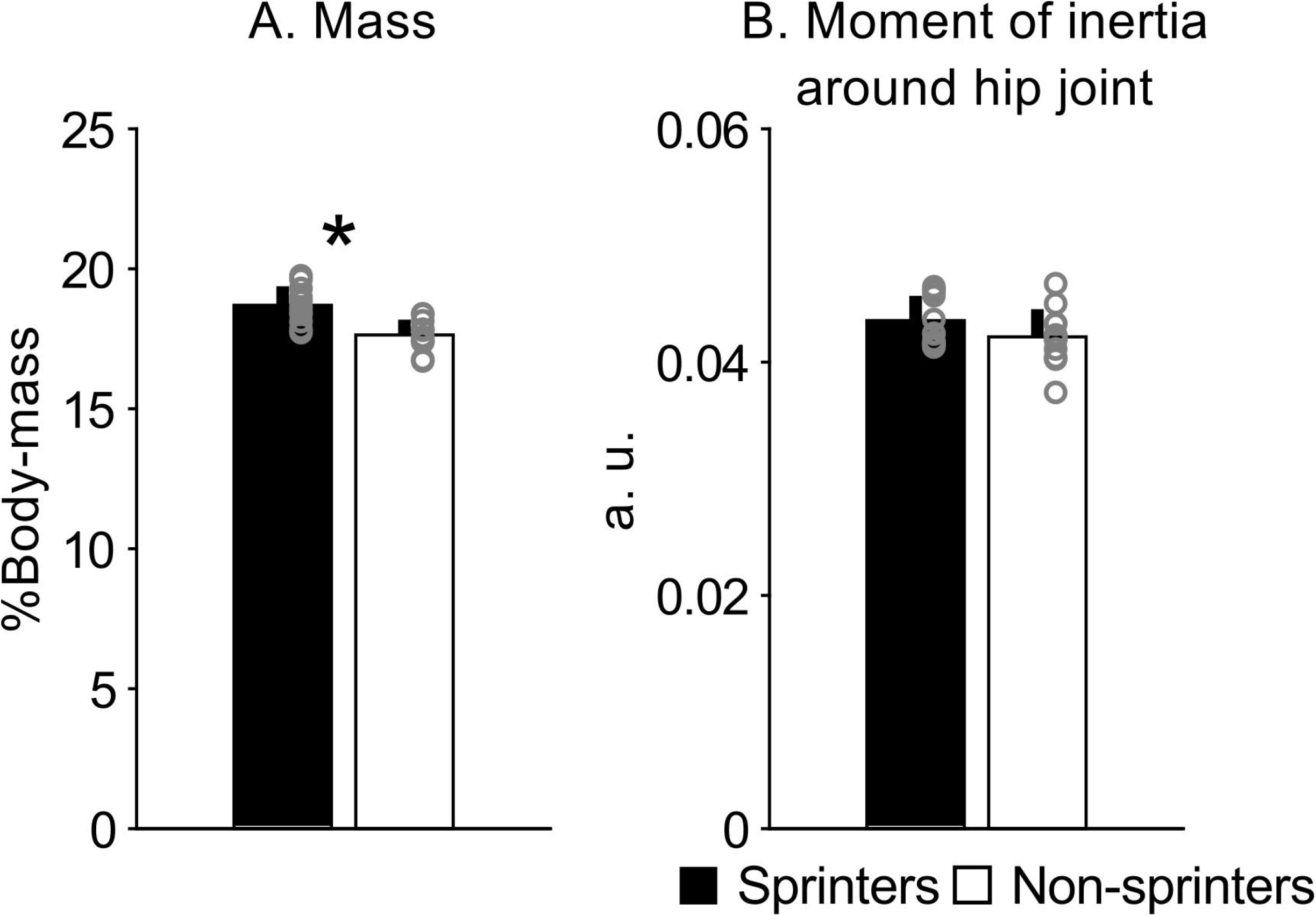
Whole lower-limb inertia properties. Relative mass (A) and normalized moment of inertia around the hip joint in hypothetically constructed anatomical position data (B). Note: Bar graphs show means and standard deviations. Gray plots show individual data. Asterisk (*) indicates a significant difference between sprinters and non-sprinters.

The relative thigh mass was significantly larger in sprinters (12.9 ± 0.4%) than in non-sprinters (11.9 ± 0.5%) (*P* < 0.01, *d* = 2.01), while those in the shank (4.5 ± 0.3% vs. 4.3 ± 0.3%) and foot (1.4 ± 0.1% vs. 1.4 ± 0.1%) were not significantly different between them (*P* = 0.13, 0.61, *d* = 0.67, 0.22; Fig. 5A). Compared to non-sprinters, all lower-limb segments in sprinters had a significantly larger lean mass ratio (thigh: 81.7 ± 2.0% vs 72.0 ± 4.8%; shank: 73.0 ± 2.0% vs 66.9 ± 3.1%; foot: 52.3 ± 2.5% vs 48.3 ± 1.9%; all *P* < 0.01, *d* = 1.85–2.62) and smaller fat mass ratio (thigh: 10.3 ± 2.1% vs. 19.2 ± 5.6%; shank: 10.2 ± 2.7% vs. 15.3 ± 3.5%; foot: 19.6 ± 1.6% vs. 24.0 ± 2.5%; all *P <* 0.01, *d* = 1.64–2.09) (Fig. 5B). The mass ratio of bone tissue in the thigh was smaller in sprinters (8.1 ± 0.6%) than in non-sprinters (8.9 ± 1.1%) (*p =* 0.03, *d* = 1.00; Fig. 5C), while those in the shank and foot were not significantly different between the groups (*P =* 0.24, 0.72, *d* = 0.51, and 0.15; Fig. 5B). The densities of all lower-limb segments were significantly larger in sprinters than in non-sprinters (thigh: 1.064 ± 0.004 g/cm^3^ vs. 1.049 ± 0.012 g/cm^3^; shank: 1.084 ± 0.007 g/cm^3^ vs. 1.073 ± 0.008 g/cm^3^; foot: 1.061 ± 0.003 g/cm^3^ vs. 1.051 ± 0.008 g/cm^3^) (all *P* < 0.01, *d* = 1.29–1.60; Fig. 5C). We showed the complete scaling coefficients to calculate the lower-limb segment inertia parameters for sprinters and non-sprinters in Table 2. The radii of gyration, the moment of inertia normalized by the segment length and segment mass, around the segmental mediolateral axis in the thigh (27.7 ± 0.3% vs 27.9 ± 0.2%; *P*=0.02, *d*=1.08) and shank (27.0 ± 0.3% vs. 27.6 ± 0.3%; *P* < 0.01, *d* = 2.24) were significantly smaller in sprinters than in non-sprinters. Although the longitudinal distance between the proximal joint and the CoM in the foot segment was significantly longer in sprinters than in non-sprinters (*P* = 0.01, *d* = 1.00), the differences in the longitudinal distances were not significant in the thigh and shank segments (*P* = 0.71, 0.15, *d* = 0.16, 0.28) (Table 2). The normalized moment of inertia of each segment of each proximal joint was not significantly different between the sprinters and non-sprinters (Table 3).

**Figure 5.**
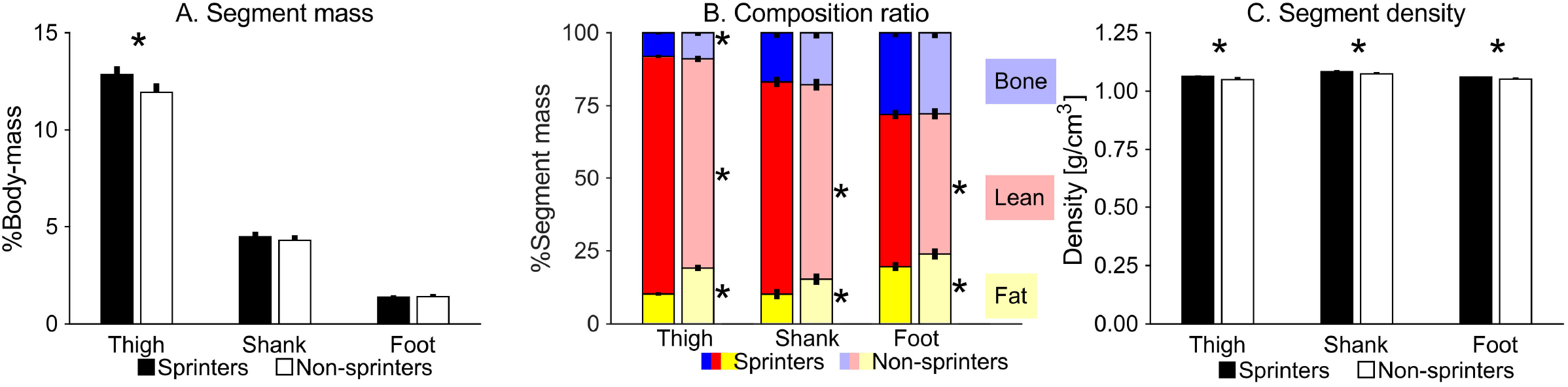
Segment mass and its determinants of each lower-limb segment. Relative mass (A), density (B), and composition ratio (C) in each of the thigh, shank, and foot segments. Note: Bar graphs show mean and standard deviation. Asterisk (*) indicates a significant difference between sprinters and non-sprinters.

**Table 2.**
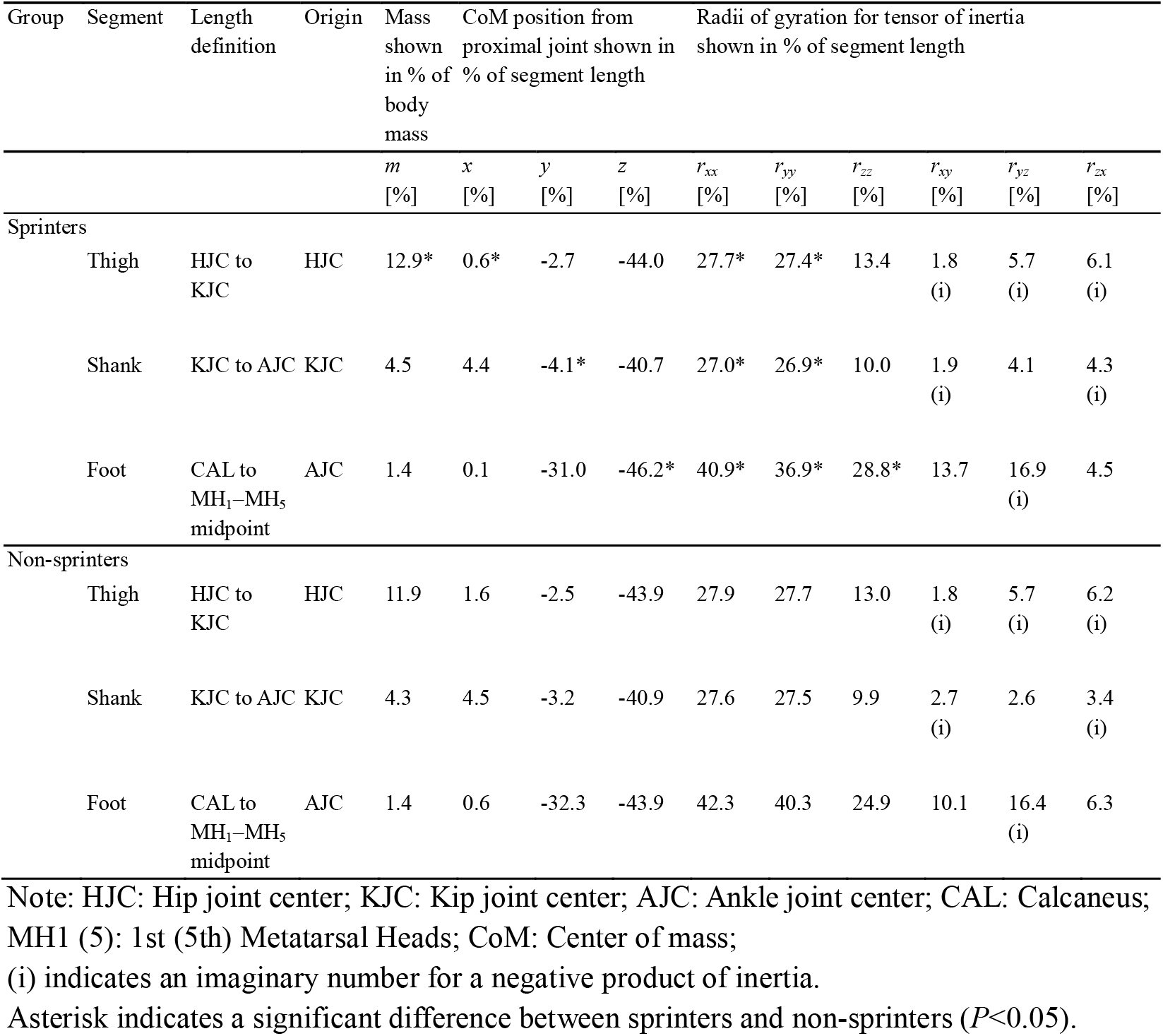
Complete segment inertia parameters in sprinters and non-sprinters.

**Table 3.** Normalized moment of inertia of each segment about the extension–flexion (or plantar flexion–dorsiflexion) axis of each proximal joint.

| Variable | Sprinters | Non-sprinters | *P*-value | Cohen’s *d* |
| --- | --- | --- | --- | --- |
| $\widehat{I}_{\text{Thigh}}^{\text{Hip}}$ [a. u.] | $0.520 \pm 0.006$ | $0.521 \pm 0.004$ | 0.80 | 0.10 |
| $\widehat{I}_{\text{Shank}}^{\text{Knee}}$ [a. u.] | $0.490 \pm 0.006$ | $0.495 \pm 0.004$ | 0.05 | 0.86 |
| $\widehat{I}_{\text{Foot}}^{\text{Ankle}}$ [a. u.] | $0.691 \pm 0.013$ | $0.691 \pm 0.014$ | >0.99 | 0.00 |

Note: $\widehat{I}_s^j$ indicates normalized moment of inertia of segment *s* (thigh, shank, or foot) about the extension–flexion (or plantar flexion–dorsiflexion) axis of the proximal joint *j* (hip, knee, or ankle).

For the values relating to the examiner’s manual processing (digitizing of the anatomical landmarks on MRIs), we confirmed the excellent repeatability in segment lengths with ICCs and CVs of 0.993 and 0.01–0.36% for the thigh, 0.994 and 0.00–0.57% for the shank, and 0.963 and 0.17–1.44% for the foot.

## Discussion

We found that sprinters have lower limbs with a larger relative mass, but not a larger 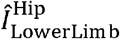 than non-sprinters. Although developed muscularity, as previously demonstrated (8–10), leads to a larger force and power exertion capacity, the increase in mass due to larger muscles has been speculated to make it difficult for the body to move in terms of inertia. However, we found that the mechanical difficulty in swinging the lower limbs in sprinters was not larger relatively small.

A notable difference in segment mass between sprinters and non-sprinters was confirmed only in the thigh (12.9% vs. 11.9%). The moment of inertia *I* is mechanically proportional to the square of the radius of gyration *r*:.

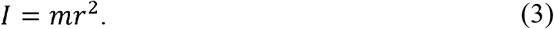

For example, if same mass (e.g., 1 kg) is added to the thigh and shank in the sprinters in this study (*r*_Hip-to-ThighCoM_: 0.19 m; *r*_Hip-to-ShankCoM_: 0.60 m), the increase in lower-limb moment of inertia due to the added mass in the thigh (0.036 kg·m^2^) is only 1/10 of that due to the added mass to the shank (0.360 kg·m^2^). Based on simple mechanical laws, a small inertia can lead to both decreasing musculoskeletal demands and increasing body segment acceleration. Thus, top-heavy, bottom-light features of the lower limb in sprinters can be an optimal solution for their performance from the perspective of inertia. The longitudinal distances between the proximal joint and the segment CoM were not significantly different between the groups. Furthermore, although the normalized moments of inertia of all segments of the proximal joints tended to be slightly smaller in sprinters, the differences were not significant. Although sprinters have inhomogeneous muscularity even within the thigh (8, 11), and differences in the mass distribution of individual muscles along the segment potentially affect inertia properties, we suggest that the moment of inertia around the hip is affected more by inter-segment differences than the mass distribution within the thigh segment.

All thigh, shank, and foot segments in sprinters had greater lean tissue ratios, smaller fat tissue ratios, and greater densities than those in non-sprinters. The sprinter/non-sprinters ratios of segment density were similar in all three segments, including the thigh (approximately 101%). The sprinter/non-sprinters ratio of thigh density was lower than that of thigh mass (107.6%). These results indicate that segment mass is more affected by segment volume than by segment density. Furthermore, the bone mass ratio of the sprinters was nearly 10% smaller than that of non-sprinters only in the thigh, which is straightforward to interpret as sprinters having a larger segment volume only in the thigh. Taken together, we indicate that the lower-limb inertia properties in sprinters, having a larger relative mass but not a larger 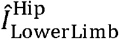, are the result of the greater volume only in the thigh.

Potential factors for the proximal-segment-specific larger mass in sprinters could be 1) training programs in sprinters, 2) inter-muscular differences in hypertrophic responses, and 3) muscle-tendon arrangement. Increasing running velocity in the high-speed range (>7 m/s) requires a larger torque/power exertion of the hip flexors and extensors (14, 15). To meet these musculoskeletal demands, sprinters may train the hip extensor and flexor muscles with particular emphasis, which could potentially be related to the proximally specific larger mass in sprinters. Meanwhile, the ankle plantar flexors exert the largest negative and positive powers during each of the early and late stance phases during sprinting (14) and play an important role in supporting the body (37). Considering such a mechanical load on the ankle plantar flexors during sprinting execution, it is unlikely that such a proximal-specific larger mass can be achieved only by daily training history. This would also be supported by our findings the trained compositional features of greater lean tissue and smaller fat tissue in sprinters than non-sprinters were confirmed in both the thigh (lean:81.7 ± 2.0% vs. 72.0 ± 4.8%, fat:10.3 ± 2.1% vs. 19.2 ± 5.6%) and shank (lean:73.0±2.0% vs. 66.9±3.1%; fat:10.2±2.7% vs. 15.3±3.5%). Meanwhile, it is relatively difficult to increase the size of the shank muscles (38), potentially due to poor protein synthesis in the shank muscles compared to other muscles (39). Even if athletes trained the thigh and shank muscles to a similar extent, the hypertrophic response to resistance training would be higher in the thigh muscles than in the shank muscles, which could also result in a proximal-specific larger mass in sprinters. Furthermore, the length ratio of the tendon to the muscle in a muscle-tendon unit (MTU) in humans has a large variation (0.01–11.25), which is generally because proximal MTUs have shorter tendons compared to distal ones (40). In a previous study (41), the Achilles tendon thickness in sprinters and non-sprinters was similar (sprinters/non-sprinters: 98.6%), whereas the ankle plantar flexor muscle thicknesses were larger in sprinters than in non-sprinters (113.8–116.8%). A previous meta-analysis (42) also showed a slight effect (weighted average effect size: 0.24) on the tendon hypertrophy in response to various types of exercise interventions. Therefore, the tendon tissue was not hypertrophic. In addition, as the tendon volume is also relatively small in comparison to the muscle volume, even if the tendon undergoes hypertrophy, the effects would be less than a similar proportion to that in the muscle. Therefore, the distal segment with a larger tendon length ratio could also contribute to the insignificant difference in the relative mass of the shank shown in this study (4.5 ± 0.3% vs. 4.3 ± 0.3%). Taken together, it is suggested that the proximal-specific larger mass in sprinters might be not just a feature of sprinters from the training strategy, but also result from an adaptation potential in humans to explosive physical exercise due to the inter-muscular differences in hypertrophic responses and the muscle-tendon arrangement. However, note that this study is a cross-sectional design with limitations to discuss adaptability in humans; longitudinal studies are needed to deepen our understanding in this regard.

We found differences in the normalized body segment inertia parameters (BSIPs), such as the radii of gyration, between sprinters and non-sprinters. Researchers have previously developed scaling coefficients to calculate BSIPs for motion analysis using direct measurements of elderly cadavers (18) and *in vivo* medical imaging such as gamma-ray scanning (16), computed tomography (20), or MRI (21, 22); however, these scaling factors do not adequately accommodate the effects of morphological characteristics in athletes on the inertial properties. We showed the complete scaling coefficients to estimate the BSIPs of the sprinter’s lower-limb segments, which provided sufficient scaling equations to calculate the hip, knee, and ankle joint dynamics. Although it is slightly outside the scope of this study, our data allowed for a more accurate analysis of motion in male sprinters.

Below, we discuss the methodological advantages and limitations of the present study. First, most of our MRI segmentation processes were computationally and automatically performed by the widely used standard Otsu’s binarization algorithm (26) on fat/water-separated images, leading to advantages in the methodological simplicity and robust results. Alternately, the landmark position coordinates were acquired by manual digitizing on the MRIs, having potential effects on the segment length used for data normalization. However, we acquired MRI images with a slice thickness of 2 mm, which is thinner than the previous studies examining inertia properties [for example, Sreenivasa et al. (27): 6 mm–48 mm; Pearsall et al. (31): 10 mm; Cheng et al. (21): 20 mm]. We confirmed excellent repeatability of the segment lengths (ICC>0.96 and CV<1.5%); thus, the potential effect of manual digitizing on the segment length would not critically alter our results. Second, non-sprinters had similar body sizes as the average Japanese 20 year old males shown in the 2019 Japanese Government statistics (43), while the sprinters were larger than the standard Japanese population, which might affect our conclusion. However, we assessed the dimensionless inertia parameters calculated according to Hof (35) to eliminate the effect of body size. Although there were significant correlations between the absolute inertia and body size parameters, the dimensionless values (the relative mass and the normalized moment of inertia of the whole lower limb around the hip joint) did not correlate with the body size, suggesting that these values could reflect the individual differences in body shapes and mass distributions, thereby not being critically affected by the inter-group differences in the body size. Third, although we determined the sample size of each group as 12 based on *a priori* power analysis (see Methods section), we excluded one of the sprinter’s data due to an error in the water/fat separation of his foot scan. However, the main findings regarding inter-group differences were statistically significant, which suggest that the exclusion did not critically affect our conclusion.

We examined only male participants. Generally, body composition has large sex differences (44); therefore, our findings might not be directly applicable to females, which is an important future theme. It is also unclear whether our findings are applicable to the upper limbs. It has been suggested that high-speed throwing in the upper limbs is one of the factors for evolution in current humans (45). It would not be surprising if humans also have morphological advantages in their upper limbs from an inertial perspective. Furthermore, unlike the lower limbs, which constantly support the body against gravity, the upper limbs, being free from such constant gravitational load, would reflect pure trainability in humans; the morphological adaptation in the upper limbs is an interesting future theme for understanding human plasticity in motor performance. Nevertheless, our findings regarding non-correspondence with an increase in lower-limb mass and moment of inertia in male sprinters open a novel perspective, inertia, for understanding human plasticity.

The present quantification of the lower-limb inertia properties in sprinters have practical implications for the training strategies. An increase in torque exertion and a smaller rate or no increase in the moment of inertia would lead to a larger angular acceleration, which could in turn facilitate improved sprint performance. In general, a trade-off relationship has been recognized between greater muscle strength and greater mechanical difficulty of moving. However, we found that the top-heavy, bottom-light feature makes the lower limb in sprinters heavier but not harder to move mechanically. This implies that sprinters can train without paying close attention to the increased mass associated with sprint-induced lower-limb muscularity and the resulting increased difficulty of moving, moment of inertia.

## Conclusion

We demonstrate the lower-limb morphology in sprinters with a larger relative mass but with a not larger normalized moment of inertia around the hip comparing it to non-sprinters. This result suggests one of the practical advantages of human morphological characteristics for explosive motor tasks, which provides a novel perspective to the literature for understanding human morphology.

## Acknowledgements

This work was supported by Yamaha Motor Foundation for Sports and JSPS KAKENHI Grant-in-Aid for Young Scientists (21K17592). We would like to thank Dr. Gaku Kakehata for his help in participant recruitment.

The results of the study are presented clearly, honestly, and without fabrication, falsification, or inappropriate data manipulation. The results of the present study do not constitute endorsement by the American College of Sports Medicine.

## TABLE LEGENDS

**Table 1 Characteristics of participants**.

**Table 2 Complete scaling coefficients to estimate segment inertia parameters in sprinters and non-sprinters**.

**Table 3 Normalized moment of inertia of each segment about the extension–flexion (or plantar flexion–dorsiflexion) axis of each proximal joint**.

## References

1. Young NM, Wagner GP, Hallgrímsson B. Development and the evolvability of human limbs. Proc Natl Acad Sci U S A. 2010;107(8):3400–5.

2. Bramble DM, Lieberman DE. Endurance running and the evolution of Homo. Nature. 2004;432(7015):345–52.

3. Lieberman DE, Bramble DM. The Evolution of Marathon Running. Sport Med. 2007;37(4):288–90.

4. Japan Sports Agency. The Report of FY 2020 Survey on Physical Strength and Athletic Performance. 2021;

5. Fukunaga T, Miyatani M, Tachi M, Kouzaki M, Kawakami Y, Kanehisa H. Muscle volume is a major determinant of joint torque in humans. Acta Physiol Scand. 2001;172(4):249–55.

6. O’Brien TD, Reeves ND, Baltzopoulos V, Jones DA, Maganaris CN. Strong relationships exist between muscle volume, joint power and whole-body external mechanical power in adults and children. Exp Physiol. 2009;94(6):731–8.

7. Balshaw TG, Maden-Wilkinson TM, Massey GJ, Folland JP. The Human Muscle Size and Strength Relationship: Effects of Architecture, Muscle Force, and Measurement Location. Med Sci Sports Exerc. 2021;53(10):2140–51.

8. Ema R, Sakaguchi M, Kawakami Y. Thigh and Psoas Major Muscularity and Its Relation to Running Mechanics in Sprinters. Med Sci Sport Exerc. 2018;50(10):2085–91.

9. Tottori N, Suga T, Miyake Y, et al. Trunk and lower limb muscularity in sprinters: what are the specific muscles for superior sprint performance? BMC Res Notes. 2021;14(1):10–5.

10. Miller R, Balshaw TG, Massey GJ, et al. The Muscle Morphology of Elite Sprint Running. Med Sci Sport Exerc. 2020;53(4):804–15.

11. Handsfield GG, Knaus KR, Fiorentino NM, Meyer CH, Hart JM, Blemker SS. Adding muscle where you need it: non-uniform hypertrophy patterns in elite sprinters. Scand J Med Sci Sport. 2017;27(10):1050–60.

12. Ema R, Wakahara T, Yanaka T, Kanehisa H, Kawakami Y. Unique muscularity in cyclists’ thigh and trunk: A cross-sectional and longitudinal study. Scand J Med Sci Sports. 2016;26(7):782–93.

13. Takahashi K, Kamibayashi K, Wakahara T. Muscle size of individual hip extensors in sprint runners: Its relation to spatiotemporal variables and sprint velocity during maximal velocity sprinting. PLoS One. 2021;16(4 April):1–12.

14. Schache AG, Blanch PD, Dorn TW, Brown NAT, Rosemond D, Pandy MG. Effect of running speed on lower limb joint kinetics. Med Sci Sports Exerc. 2011;43(7):1260–71.

15. Dorn TW, Schache AG, Pandy MG. Muscular strategy shift in human running: dependence of running speed on hip and ankle muscle performance. J Exp Biol. 2012;215:1944–56.

16. Huang HK, Wu SC. The evaluation of mass densities of the human body in vivi from CT Scans. Comput Biol Med. 1976;6(4):337–43.

17. Martin PE, Mungiole M, Maezke MW, Longhill JM. The use of magnetic resonance imaging for measuring segment inertial propertierties. J Biomech. 1989;22(4):367–76.

18. Clauser CE, McConville JT, Young JW. Weight, volume, and center of mass of segments of the human body (AMRL-TR-69-70). Wright-Patterson Air Force Base, Ohio. 1969;1–112.

19. de Leva P. Adjustments to Zatsiorsky-Seluyanov’s segment inertia parameters. J Biomech. 1996;29:1223–30.

20. Huang HK, Suarez FR. Evaluation of cross-sectional geometry and mass density distributions of humans and laboratory animals using computerized tomography. J Biomech. 1983;16(10):821–32.

21. Cheng CK, Chen HH, Chen CS, Lee CL, Chen CY. Segment inertial properties of Chinese adults determined from magnetic resonance imaging. Clin Biomech. 2000;15(8):559–66.

22. Mungiole M, Martin PE. Estimating segment inertial properties: Comparison of magnetic resonance imaging with existing methods. J Biomech. 1990;23(10):1039–46.

23. Shan G, Bohn C. Anthropometrical data and coefficients of regression related to gender and race. Appl Ergon. 2003;34(4):327–37.

24. Sawilowsky SS. New Effect Size Rules of Thumb. J Mod Appl Stat Methods. 2009;8(2):597–9.

25. Klingensmith JD, Elliott AL, Fernandez-del-Valle M, Mitra S. Automated segmentation of cardiac adipose tissue in Dixon magnetic resonance images. J Biomed Graph Comput. 2017;8(1):1.

26. Otsu N. A threshold selection method from gray-level histograms. IEEE Trans Syst Man Cybern. 1996;9(1):62–6.

27. Sreenivasa M, Chamorro CJG, Gonzalez-Alvarado D, Rettig O, Wolf SI. Patient-specific bone geometry and segment inertia from MRI images for model-based analysis of pathological gait. J Biomech. 2016;49(9):1918–25.

28. Harrington ME, Zavatsky AB, Lawson SEM, Yuan Z, Theologis TN. Prediction of the hip joint centre in adults, children, and patients with cerebral palsy based on magnetic resonance imaging. J Biomech. 2007;40(3):595–602.

29. Rossi M, Lyttle A, El-Sallam A, Benjanuvatra N, Blanksby B. Body segment inertial parameters of elite swimmers using DXA and indirect methods. J Sport Sci Med. 2013;12(4):761–75.

30. Pearsall DJ, Reid JG, Livingston L a. Segmental inertial parameters of the human trunk as determined from computed tomography. Ann Biomed Eng. 1996;24(2):198–210.

31. Pearsall DJ, Reid JG, Ross R. Inertial properties of the human trunk of males determined from magnetic resonance imaging. Ann Biomed Eng. 1994;22(6):692–706.

32. Sado N, Yoshioka S, Fukashiro S. A non-orthogonal joint coordinate system for the calculation of anatomically practical joint torque power in three-dimensional hip joint motion. Int J Sport Heal Sci. 2017;15:111–9.

33. Sado N, Yoshioka S, Fukashiro S. Curved Approach in High Jump Induces Greater Jumping Height without Greater Joint Kinetic Exertions than Straight Approach. Med Sci Sports Exerc. 2022;54(1):120–8.

34. Dumas R, Chèze L, Verriest JP. Adjustments to McConville et al. and Young et al. body segment inertial parameters. J Biomech. 2007;40(3):543–53.

35. Hof AL. Scaling gait data to body size. Gait Posture. 1996;4(3):222–3.

36. Cohen J. Statistical power analysis for the behavioral sciences. 2nd ed. Hillsdale: Lawrence Earlbaum Associates; 1988.

37. Pandy MG, Lai AKMM, Schache AG, Lin Y-CC. How muscles maximize performance in accelerated sprinting. Scand J Med Sci Sports. 2021;31(July):1882–96.

38. Weiss LW, Clark FC, Howard DG. Effects of heavy-resistance triceps surae muscle training on strength and muscularity of men and women. Phys Ther. 1988;68(2):208–13.

39. Trappe TA, Raue U, Tesch PA. Human soleus muscle protein synthesis following resistance exercise. Acta Physiol Scand. 2004;182(2):189–96.

40. Hoy MG, Zajac FE, Gordon ME. A musculoskeletal model of the human lower extremity: The effect of muscle, tendon, and moment arm on the moment-angle relationship of musculotendon actuators at the hip, knee, and ankle. J Biomech. 1990;23(2):157–69.

41. Kubo K, Ikebukuro T, Yata H, Tomita M, Okada M. Morphological and mechanical properties of muscle and tendon in highly trained sprinters. J Appl Biomech. 2011;27(4):336–44.

42. Bohm S, Mersmann F, Arampatzis A. Human tendon adaptation in response to mechanical loading: a systematic review and meta-analysis of exercise intervention studies on healthy adults. Sport Med - Open [Internet]. 2015;1(1) doi:10.1186/s40798-015-0009-9.

43. Government of Japan. e-Stat: Portal Site of Official Statistics of Japan. 2020; Available from: https://www.e-stat.go.jp/en.

44. Bredella MA. Sex Differences in Body Composition BT - Sex and Gender Factors Affecting Metabolic Homeostasis, Diabetes and Obesity. In: Mauvais-Jarvis F, editor. Cham: Springer International Publishing; 2017. p. 9–27. Available from: https://doi.org/10.1007/978-3-319-70178-3_2.

45. Roach NT, Venkadesan M, Rainbow MJ, Lieberman DE. Elastic energy storage in the shoulder and the evolution of high-speed throwing in Homo. Nature. 2013;498(7455):483–6.

