## Supplemental Table for "Lower limb in sprinters with larger relative mass but not larger normalized moment of inertia"

| **Supplemental Table S1. Selected dicom meta data.** | |
| --- | --- |
| Variable Name | Value |
| Format | DICOM |
| FormatVersion | 3 |
| Width | 1024 |
| Height | 1024 |
| BitDepth | 16 |
| ColorType | grayscale |
| SpecificCharacterSet | ISO_IR100 |
| ImageType | DERIVED\PRIMARY\DIXON\FAT |
| Manufacturer | GEMEDICALSYSTEMS |
| SeriesDescription | FAT 3D Ax LAVA Flex Thigh |
| ManufacturerModelName | SIGNAPremier |
| VolumetricProperties | VOLUME |
| ScanningSequence | GR |
| MRAcquisitionType | 3D |
| SliceThickness | 4 |
| RepetitionTime | 4.344 |
| EchoTime | 1.732 |
| NumberOfAverages | 1.5805 |
| ImagingFrequency | 127.8062 |
| EchoNumber | 1 |
| MagneticFieldStrength | 3 |
| SpacingBetweenSlices | 2 |
| EchoTrainLength | 1 |
| PercentSampling | 100 |
| PercentPhaseFieldOfView | 100 |
| PixelBandwidth | 325.527 |
| ReconstructionDiameter | 300 |
| ReceiveCoilName | 30AA+60PA |
| InPlanePhaseEncodingDirection | COL |
| FlipAngle | 12 |
| VariableFlipAngleFlag | N |
| SAR | 1.6502 |
| PatientPosition | FFS |
| SamplesPerPixel | 1 |
| PhotometricInterpretation | MONOCHROME2 |
| Rows | 1024 |
| Columns | 1024 |
| PixelSpacing | 0.2930 0.2930 |
| BitsAllocated | 16 |
| BitsStored | 16 |
| HighBit | 15 |

Supplemental Table S2. Anatomical landmarks to be acquired by manual digitizing.

| Segments | Landmarks |
| --- | --- |
| Thigh | Surface of the femoral head (30 points) |
|  | Lateral articular cleft between the femur and tibia condyle |
|  | Medial articular cleft between the femur and tibia condyle |
| Shank | Lateral articular cleft between the femur and tibia condyle |
|  | Medial articular cleft between the femur and tibia condyle |
|  | Malleolus lateralis |
|  | Malleolus medialis |
| Foot | Malleolus lateralis |
|  | Malleolus medialis |
|  | Calcaneus |
|  | 1st meta-tarsal heads |
|  | 5th meta-tarsal heads |
|  | 2nd toe tip |

Note: Anatomical landmarks other than the femoral head were digitized closest points on the skin (i.e., points to which reflective markers can be attached when doing motion capture).
